# CCA based multi-view feature selection for multi-omics data integration

**DOI:** 10.1101/243733

**Authors:** Yasser El-Manzalawy

## Abstract

Recent technological advances in high-throughput omics technologies and their applications in genomic medicine have opened up outstanding opportunities for individualized medicine. However, several challenges arise in the integrative analysis of such data including heterogeneity and high dimensionality of the omics data. In this study, we present a novel multi-view feature selection algorithm based on the well-known canonical correlation analysis (CCA) statistical method for jointly selecting discriminative features from multi-omics data sources (multi-views). Our results demonstrate that models for predicting kidney renal clear cell carcinoma (KIRC) survival using our proposed method for jointly selecting discriminative features from copy number alteration (CNA), gene expression RNA-Seq, and reverse-phase protein arrays (RPPA) views outperform models trained using single-view data as well as three integrated models developed using data fusion approaches including CCA-based feature fusion.

## I. Introduction

Translational bioinformatics, an emerging field in the study of health informatics, has become a key component in biomedical research and an important discipline in the era of precision medicine [1]. Advances in high-throughput omics technologies (genomics, transcriptomics, proteomics, and metabolomics) have provided new opportunities for integrated and data-intensive analyses of omics data to deciphering genotype-phenotype interactions in complex diseases such as cancer [2].

A major challenge in translational bioinformatics is developing models for predicting clinical outcome using multiomics data for improved diagnostics, prognostics, and further therapeutics [3]. Amongst different clinical outcome prediction tasks, predicting cancer survival using omics profiles is a major challenge in translational bioinformatics [4]. In general, existing approaches for developing cancer survival models from multi-omics data can be categorized as either *data fusion* approaches, where multiple data sources are combined to form a single dataset, or *model fusion* approaches, where models trained using independent omics data sources are combined into a single consensus or meta-model. Alternatively, an emerging machine learning research direction, called multi-view learning [5], attempts to develop models that learn (or select features) jointly from multiple data sources (i.e., multiple views) and thus improve the generalization performance of the learned models.

Multi-view learning algorithms attempt to learn one model from each view while jointly optimizing these view-specific models to improve the generalization performance. Recently, several supervised and unsupervised multi-view learning algorithms have been proposed including multi-view support vector machines [6], multi-view boosting [7], multi-view *k*-means [8], and clustering via canonical correlation analysis [9]. Besides supervised and unsupervised learning problems, several traditional single-view machine learning problems have been extended to the multi-view settings such as multi-view semi-supervised learning [10-12], multi-view transfer learning [13], multi-view feature extraction [14], and multi-view dimensionality reduction [15, 16]. A good representative example of the multi-view dimensionality reduction algorithms is the Canonical Correlation Analysis (CCA) algorithm [17] which is an early statistical method for reducing the dimensionality of a pair of datasets. CCA jointly learns a shared subspace between two datasets (i.e., views) and is, therefore, considered as one of the early multi-view learning algorithms. Several variants of CCA have been proposed for further extending the classical CCA to more than two views and to learn non-linear subspaces [18].

Against this background, we believe that multi-view learning is a promising machine learning direction for tackling arising challenges in data integration. However, the vast majority of existing multi-view learning algorithms are not designed to effectively learn intrinsic relationships across high-dimensional views [14]. To address this limitation, we utilize multi-view feature selection for the integration of high-dimensional multi-omics data. Specifically, we present a novel multi-view feature selection algorithm that uses the CCA learned projective transformation to score and rank input features from multiple views (i.e., omics data sources). We compare our approach against two baseline data fusion approaches as well as against an approach for serial feature fusion [13] applied to sets of features in the CCA reduced dimensionality space [19]. Experimental results on predicting kidney renal clear cell carcinoma (KIRC) survival [20, 21] (using copy number alteration (CNA), gene expression RNA-Seq, and reverse-phase protein arrays (RPPA) as three input views) demonstrate superior performance of our proposed method.

## II Materials and Methods

### A. Datasets

Clinical data as well as normalized and preprocessed copy number alteration (CNA), gene expression RNA-Seq, and reverse-phase protein arrays (RPPA) data of the kidney renal clear cell carcinoma (KIRC) TCGA cohort were downloaded from UCSC Xena functional genomics browser [22]. Table I overviews the KIRC dataset. Since CCA is applicable to only complete views, only patients with CNA, RNA-Seq, RPPA, and survival information were kept. Instances corresponding to patients with survival time ≥ 3 years were labeled as long-term survivors (positive instances) while patients with survival time < 3 years and vital status of 0 were labeled as short-term survivors (negative instances). Thus, the final dataset contains 256 and 103 positive and negative samples, respectively. Each final data source was then pre-filtered and normalized as follows: i) feature values in each sample were re-scaled to lie in the interval [0,1]; ii) features with variance less than 0.02 were removed.

**TABLE I.**
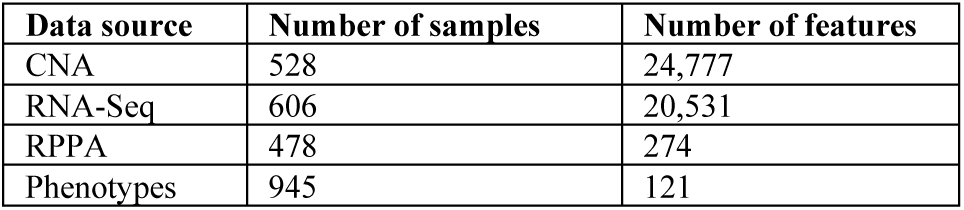
Overview of KIRC dataset

### B. Canonical Correlation Analysis (CCA)

We start with convenient notations that will be used throughout the present work (See Table II).

Canonical correlation analysis [17] is a commonly used statistical method for finding correlation relationships between two sets of features (i.e., two views or representations of the same set of observations). Given two zero-mean unlabeled datasets 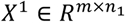 and 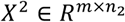, CCA determines two projective transforms 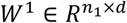 and 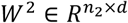 that maximize the linear correlation between the projections of the two datasets in the *d*-dimensional space. For more details about the derivation and solution of CCA, please see [23].

**TABLE II.**
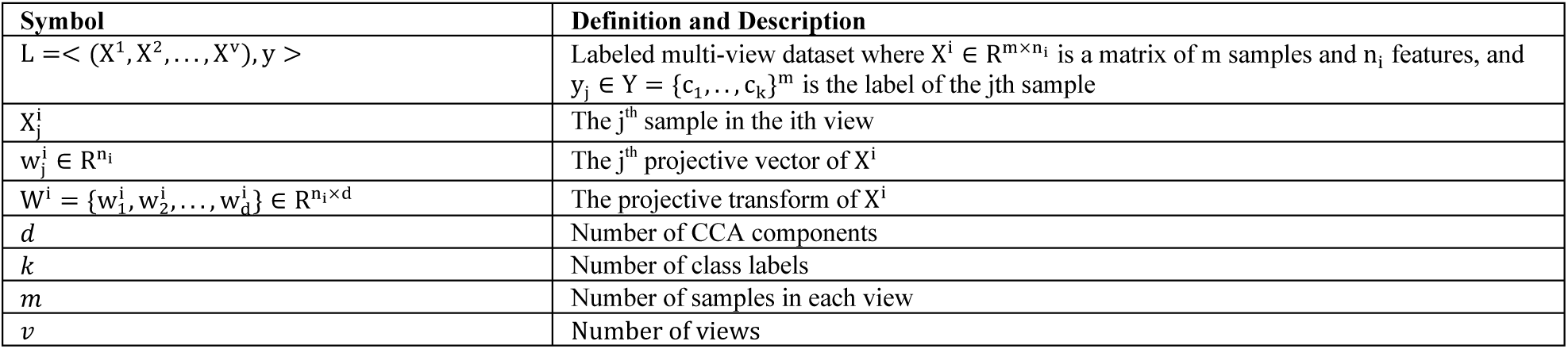
Notations

| Symbol | Definition and Description |
| --- | --- |
| $L = \langle (X^1, X^2, \dots, X^v), y \rangle$ | Labeled multi-view dataset where $X^i \in \mathbb{R}^{m \times n_i}$ is a matrix of $m$ samples and $n_i$ features, and $y_j \in Y = \{c_1, \dots, c_k\}^m$ is the label of the $j$th sample |
| $X_j^i$ | The $j$th sample in the $i$th view |
| $w_j^i \in \mathbb{R}^{n_i}$ | The $j$th projective vector of $X^i$ |
| $W^i = \{w_1^i, w_2^i, \dots, w_d^i\} \in \mathbb{R}^{n_i \times d}$ | The projective transform of $X^i$ |
| $d$ | Number of CCA components |
| $k$ | Number of class labels |
| $m$ | Number of samples in each view |
| $v$ | Number of views |

In this work, we used a recently developed Python library for Regularized Kernel Canonical Correlation Analysis (Pyrcca) [18]. Unlike other existing CCA implementations, Pyrcca provides support for L2 regularization as well as kernalized CCA. Moreover, Pyrcca implements an extension of the generalized eigenvalue problem [24] to support the applicability of CCA to more than two datasets.

### C. CCA-based Feature Fusion (CCAFF)

Sun et al. [19] presented a CCA-based feature fusion method for mapping two labeled views into a single labeled dataset in *2d*-dimentional space. Here, we generalize their method to more than two views.

Given a labeled multi-view dataset L = < (X^1^, X^2^, …, X^v^), y >, our implementation of the CCA-based feature fusion (CCAFF) applies Pyrcca to the unlabeled multi-view data < (X^1^, X^2^, …, X^v^), y > to obtain the projective transformations W^1^, W^2^,…, W^*v*^ corresponding to *d* generalized eigenvalues. Then, the serial feature fusion strategy [13] is applied to fuse the input labeled multi-view data into a labeled single-view dataset < X^fused^, y > in *vd*-dimensional space where the linear correlation between projected views is maximized and X^fused^ is the concatenation of (W^1^) ^T^ X^1^, (W^2^) ^T^ X^2^,…, and (W^v^) ^T^ X^v^.

### D. CCA-based Multi-view Feature Selection (CCAFS)

Assuming there is a good common *d*-dimensional space among all views in an unlabeled multi-view data < (X^1^, X^2^, …, X^v^), y > that could be obtained using the CCA method, we propose to use the learned projective transformation matrices to score and rank the input features in each view and to use the concatenation of top selected features from each view as an integrated representation of the input views. This approach could be viewed as a multi-view feature selection approach since the features are jointly selected from the multiple views based on CCA determined projective transformations.

In Artificial Neural Network (ANN) literature, there are different methods for quantifying the contributions of input variables in ANN models (see [25] for an interesting comparison of such methods). In this work, we adapted the weights method [25] to determine the contribution of each input feature using the CCA projective transformation matrix. Specifically, given the *i*^th^ projective transformation matrix 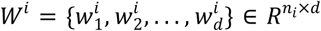, the score of each input feature *j* for *j* = 1, …, *n*_*i*_ was computed as follows:

1) Compute normalized weights matrix, *Q*^*i*^, for the *i*^th^ view by normalizing the absolute value of each element 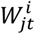 by the sum of the absolute values of the *t*^*t*h^ column, i.e.

~~~
for *t* = 1 to *d*
   for *j* = 1 to *n*_*i*_
~~~

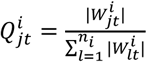

~~~
   end
end
~~~

2) Determine the importance score of the *j*^th^ feature in the *i*^th^ view using Eq. 1

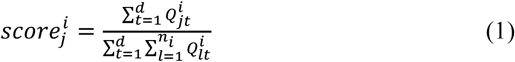

Algorithm 1 summarizes our proposed CCA-based feature selection (CCAFS) algorithm. The inputs to the algorithm are: labeled multi-view dataset, *L*; number of components, *d*; number of features to select, *k*; value of regularization parameter for CCA method, *λ*; a weight vector controlling how many features to be selected from each view, *w*.

#### Algorithm 1. CCAFS

**Requires**: L = >(X^1^,X^2^,…,X^v^),*y*>, *d,k,λ,w* = [w_1_, …, w_v_]

1: num_*i*_ = ⌊ *w*_*i*_*k*⌋ for i=1,2,3,… κ

2: W^1^,W^2^,…,W^v^ ← Pyrcca([X^1^, X^2^, …, X^v^],*d,λ*)

3 Compute the normalized weights matrices, *Q*^*i*^’s

4: Compute features importance scores using Eq. 1

5: Return the indices of top num_*i*_ features for *i* = 1,2, …, *ν*

### E. Experimental Settings

We experimented with three broadly used machine learning algorithms: Random Forest (RF) [26], eXtreme Gradient Boosting (XGB) [27], and Logistic Regression (LR) [28]. These three algorithms are implemented in sklearn machine learning library [29]. Unless stated otherwise, we used the default parameters of these algorithms except for the number of estimators (i.e., number of tree classifiers for RF and XGB models) where we set it to 500.

For feature selection methods applied to single-view datasets, we used an embedded filter [30] based on an RF classifier trained using 500 trees. We also experimented with a Two-Stage (TS) filter selection method by applying the RF-based filter twice in order to select *k* ≤ 100 features such that the first filter selects 100 features from the input features while the second filter selects *k* features from the 100 features selected in the first stage.

For data integration methods, we applied the Two-Stage data fusion strategy summarized in Fig. 1. Briefly, at the first stage, RF-based filters were used to select top 100 features from each individual view. The second stage represents the data fusion step where either CCAFF or CCAFS filter was used to integrate the 100 features from each view into integrated/fused features. If a RF-based filter was used in the second stage, this filter would be applied to the concatenation of Stage II inputs.

**Figure 1.**
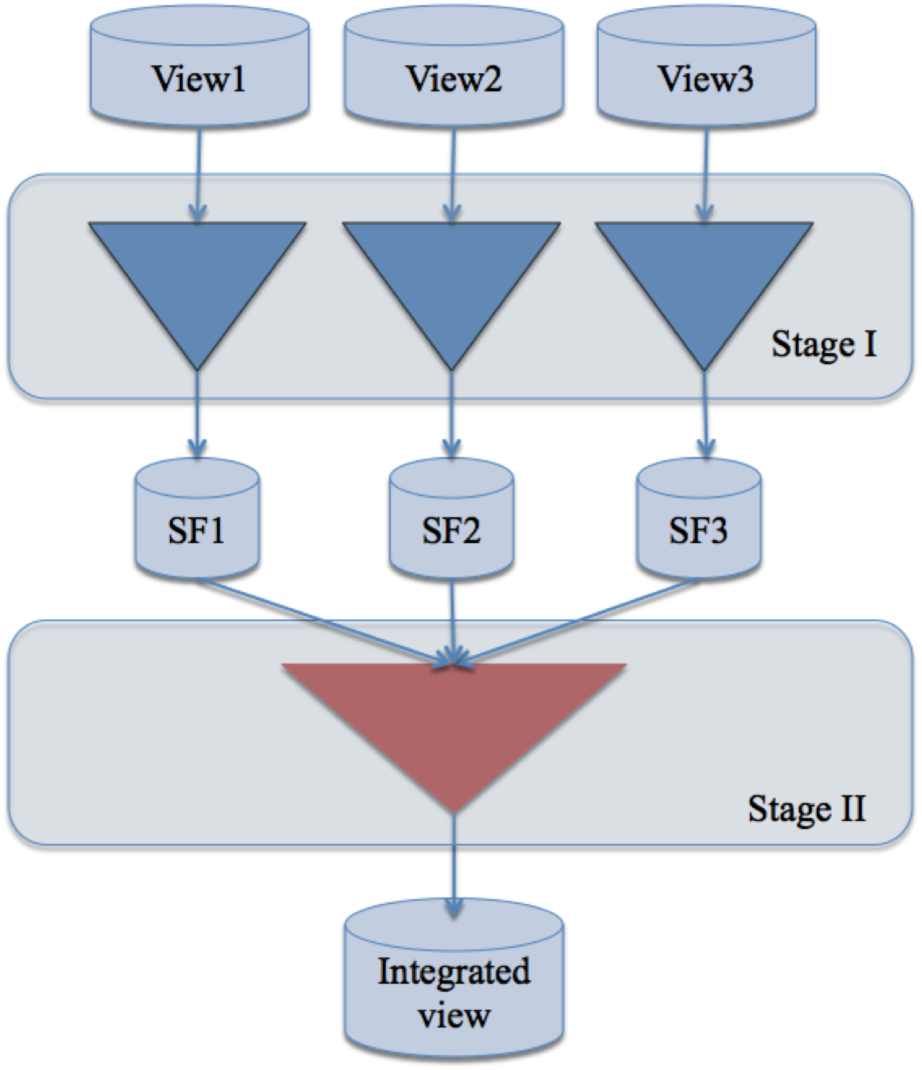
Overview of the Two-Stage framework for integrating KIRC multi-omics data.

In the case of CCAFF and CCAFS methods, we set the Pyrcca regularization parameter to 0.1 and experimented with *d* = 10, 20, …, 50 canonical componenets. For CCAFS, we set the weights to 0.4, 0.3, and 0.3 for RNA-Seq, RPPA, and CNA views, respectively. Our choice of these weights was based on the performance of view-specific models (see Section III.A).

We used the 5-fold cross-validation procedure to evaluate our classifiers and we assessed their predictive performance using the following widely used threshold-dependent and threshold-independent metrics [31]: Accuracy (ACC), Sensitivity (Sn), Specificity (Sp), Matthews Correlation Coefficient (MCC), and Area Under ROC Curve (AUC). It should be noted that feature selection methods were applied at each iteration in the 5-fold cross-validation procedure such that selected features are determined using training data only.

## III. Results and Discussion

### A. Performance Comparisons of View-specific Models

Our first set of experiments was conducted to determine which individual data source (view) could be used to build good prediction models of KIRC survival. Fig. 2 shows the AUC of different classifiers evaluated using 5-fold cross-validation and the top *k* = {10, 20, 30,…, 100} features selected using the feature importance scores inferred from a RF classifier trained using training data at each fold. Interestingly, classifiers developed using CNA seem to have poor classification performance with the highest observed AUC equals 0.59. On the other hand, the vast majority of the classifiers developed using RNA-Seq data have substantially better performance (e.g., AUC close to 0.70). The best performing RNA-Seq based classifier is RF trained and tested using only top 10 selected features. Finally, classifiers using RPPA data have AUC centered around 0.65 and the highest AUC of 0.69 is obtained using LR and top 20 selected features.

**Figure 2.**
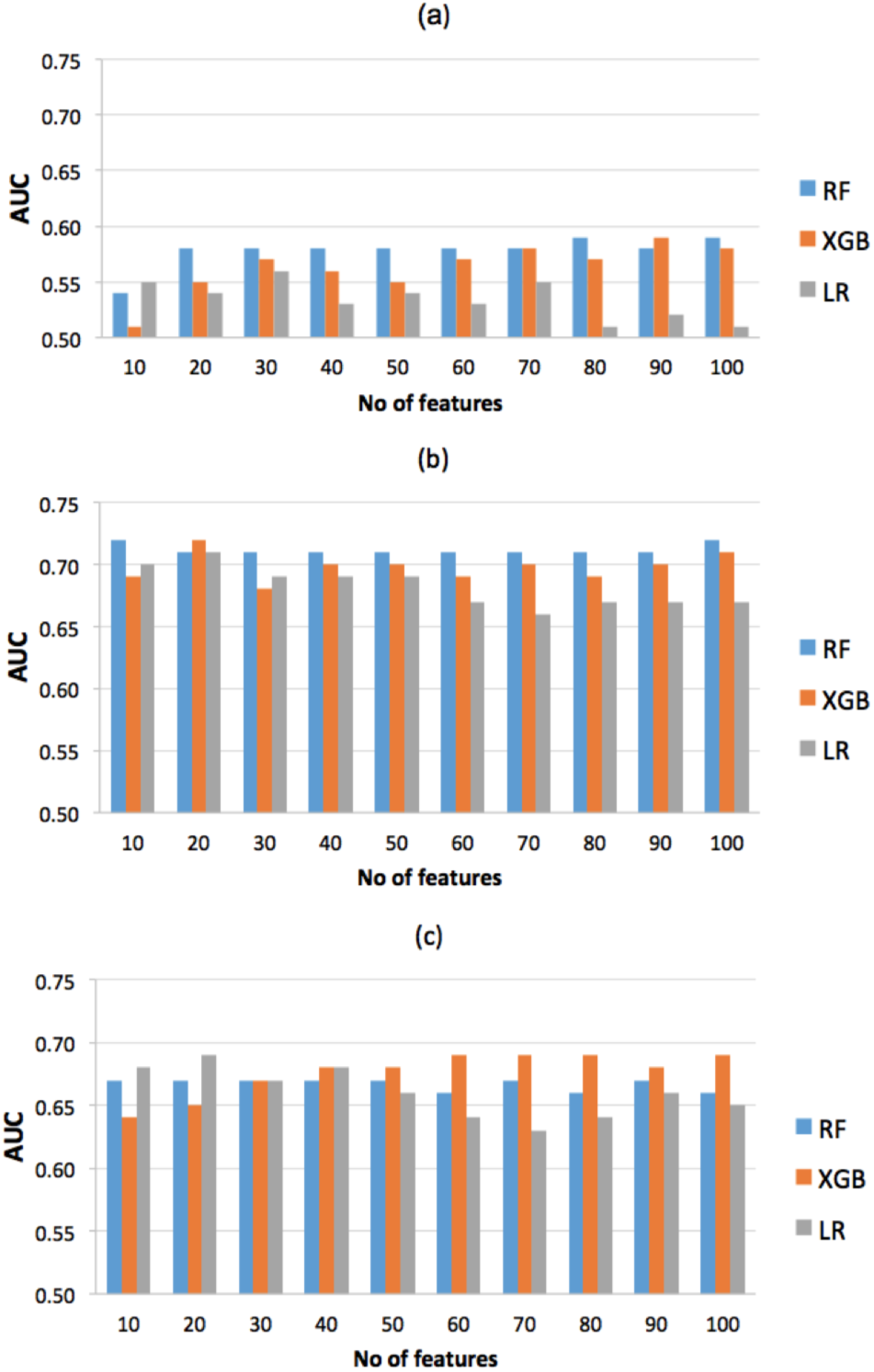
AUC scores for view-specific models evaluated using a) CNA, b) RNA-Seq, and c) RPPA views of the KIRC dataset.

Unsurprisingly, our results indicate that both the best machine learning algorithm and the optimal number of features for predicting KIRC survival are view-specific. For CNA and RNA-Seq views, RF models consistently outperformed XGB and LR models for the vast majority of choices of the number of features, while for RPPA view, RF models were outperformed by either XGB or LR models for all choices of *k* except for *k* = 30.

Our choice of embedded feature selection method using a RF classifier was based on exploratory analysis and evaluation of several feature selection methods including RF feature importance [26]; Lasso [32]; ElasticNet [33]; and Recursive Feature Elimination (RFE) [34] (data not shown). Results of this analysis also suggested that a Two-Stage (TS) feature selection approach is better than a Single-Stage (SS) feature selection. In this Two-Stage feature selection approach, an embedded feature selection method using RF classifier was used to select the top 100 features and then a second RF-based filter was used to select the top *k* features out of the top 100 features. Table III compares the top performing view-specific classifiers obtained using SS and TS feature selection methods. For CNA data, the best RF classifier using top 10 TS features has 0.03 improvement in AUC compared to the best RF classifier using top 100 SS features. The improvement is substantial when we compare RF classifiers trained using SS and TS selected features. Using SS selected features, the best performing RF classifier uses 100 features and has an AUC of 0.54 whereas using TS selected features, a higher AUC of 0.61 is observed using only 10 selected features. For RNA-Seq data, the best performing classifier using TS feature selection method has 0.01 improvement in AUC compared to the best performing classifier using SS feature selection method. Surprisingly, for RPPA dataset, slightly better AUC is observed using SS feature selection method. One possible justification is that the RPPA dataset has only 274 features and Single-Stage feature selection is sufficient. Based on these results, our data fusion models, reported in the following subsections, used the Two-Stage feature selection framework in Fig. 1.

**TABLE III.** Performance comparison of top performing view-specific models using single-stage (ss) and two-stage(ts) feature selection.

| Method | View | Classifier | No of features | ACC | Sn | Sp | MCC | AUC |
| --- | --- | --- | --- | --- | --- | --- | --- | --- |
| SS | CNA | RF | 100 | 0.71 | 0.91 | 0.20 | 0.16 | 0.59 |
| SS | RNA-Seq | RF | 10 | 0.74 | 0.88 | 0.39 | 0.30 | 0.72 |
| SS | RPPA | LR | 20 | 0.62 | 0.69 | 0.45 | 0.15 | 0.69 |
| TS | CNA | RF | 10 | 0.71 | 0.92 | 0.19 | 0.17 | 0.62 |
| TS | RNA-Seq | LR | 10 | 0.71 | 0.82 | 0.41 | 0.27 | 0.73 |
| TS | RPPA | RF | 20 | 0.74 | 0.92 | 0.29 | 0.28 | 0.68 |

### B. Performance Comparisons of Baseline Data Fusion Models

We report the results of two baseline data fusion approaches (both were applied to the top 100 features selected from each view using RF-based embedded filter). The first approach, called Concatenation of Input Features (CIF), concatenates the 100 input features from each view into a single combined view and then uses a second stage RF-based embedded filter for selecting the top *k* features. The major limitation of this basic approach is that it does not keep track of the source (i.e. view) of each feature. The second basic approach is called Concatenation of Selected Features (CSF). Briefly, CSF uses a second stage RF-based embedded filter for selecting the top ⌊*k*/*v*⌋ features out of the 100 input features selected at first stage from each view, where *v* is the number of views. The major limitation of CSF is that each view is treated independently from other views and thus it ignores complementary relationships between features in different views.

Fig. 3 shows the AUC scores for different classifiers constructed using CIF and CSF data fusion approaches for integrating CNA, RNA-Seq, and RPPA views. For models evaluated using the CIF data fusion approach, XGB classifiers outperform RF and LR classifiers for *k* ≥ 50 and the highest observed AUC is 0.74 for *k* = 60 or 70. For models evaluated using the CSF approach, RF classifiers have the highest AUC score for all values of *k* considered in our experiments except for *k* = 20 and 30 where LR classifiers have the highest AUC scores of 0.71 and 0.74, respectively.

**Figure 3.**
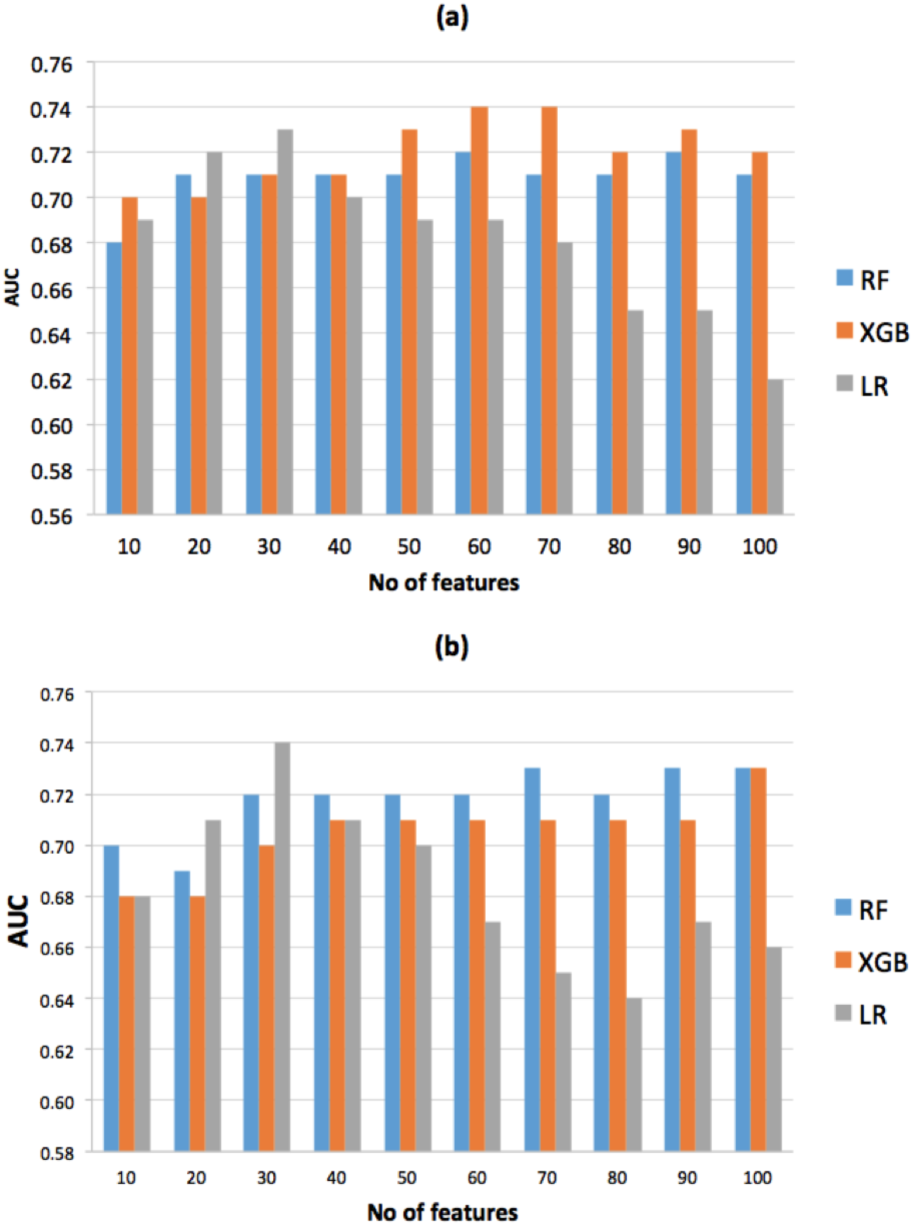
AUC scores for different classifiers developed using two baseline data fusion methods: a) Concatenation of Input Features (CIF), b) Concatenation of Selected Features (CSF).

So far, the best performing models for predicting KIRC survival were obtained using LR algorithm and either top 10 RNA-Seq features (AUC = 0.73) or top 30 integrated features selected using the CSF method (AUC = 0.74). Next, we report the predictive performance of multi-view models based on CCA feature fusion [19] and our proposed CCA feature selection algorithm.

### C. Multi-view Models Outperform View-specific and Baseline Data Fusion Models

We evaluated CCA feature selection (CCAFS) and CCA feature fusion (CCAFF) models for integrating CNA, RNA-Seq, and RPPA views by projecting the top 100 features selected from each view into a common *d*-dimensional space where *d* = 10, 20,…, 50 using the CCA algorithm. Due to space limitation, we report only the best results obtained using *d* = 50.

Fig. 4 reports the AUC scores for different classifiers evaluated using the top *k* features selected jointly from the three views using our proposed CCAFS method. We note that the XGB classifier outperforms RF and LR classifiers for k ≥ 30 and the highest reachable AUC score of 0.76 is reported using *k* = 100. This XGB classifier outperforms the best view-specific model (AUC = 0.73) as well as the best CIF and CSF models (AUC = 0.74). Moreover, this XGB classifier has 0.05 increase in AUC when compared to the best view-specific XGB classifier (see Fig. 2).

**Figure 4.**
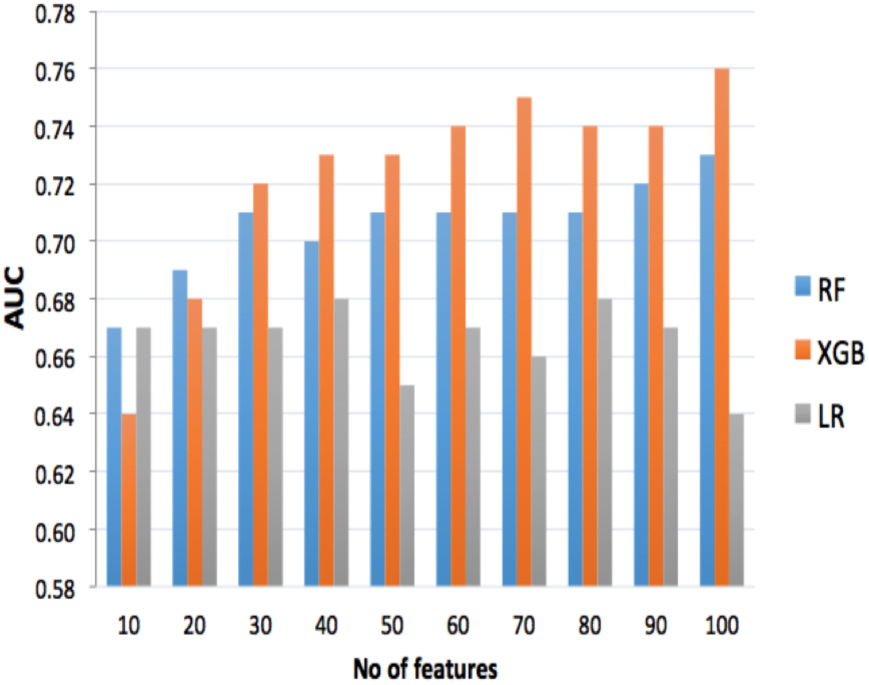
AUC scores of three classifiers evaluated using CCA-based Feature Selection (CCAFS)

Table IV compares the top performing RF, XGB, and LR classifiers built using CCAFS and CCAFF multi-view features. Better performance metrics are noted for RF and XGB models trained using CCAFS based integrated views while for LR classifiers, better performance measures are observed using the CCAFF approach. A major limitation of CCAFF models is that these models are hard to interpret since the input to the classifier is the projected views. Fortunately, CCAFS based models are trained using features selected jointly from the multiple views and, therefore, enables the examination of the learned models for getting insights and for identifying key features (e.g., biomarkers for KIRC survival).

**TABLE IV.**
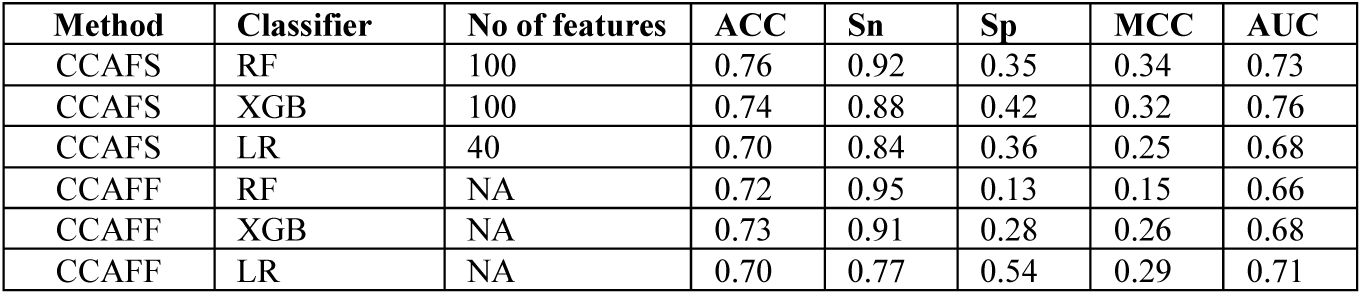
Performance comparison of CCAFS and CCAFF models using three classifiers for predicting kirc survival using three multi-omics views.

| Method | Classifier | No of features | ACC | Sn | Sp | MCC | AUC |
| --- | --- | --- | --- | --- | --- | --- | --- |
| CCAFS | RF | 100 | 0.76 | 0.92 | 0.35 | 0.34 | 0.73 |
| CCAFS | XGB | 100 | 0.74 | 0.88 | 0.42 | 0.32 | 0.76 |
| CCAFS | LR | 40 | 0.70 | 0.84 | 0.36 | 0.25 | 0.68 |
| CCAFF | RF | NA | 0.72 | 0.95 | 0.13 | 0.15 | 0.66 |
| CCAFF | XGB | NA | 0.73 | 0.91 | 0.28 | 0.26 | 0.68 |
| CCAFF | LR | NA | 0.70 | 0.77 | 0.54 | 0.29 | 0.71 |

It is worth noting that all the models summarized in Table IV have low specificity. Fortunately, higher specificities could be obtained by increasing the threshold for converting predicted probabilities into binary labels. For estimating threshold-dependent metrics in Table IV, we used an arbitrary threshold of 0.5 such that instances with predicted probabilities greater than 0.5 were assigned positive labels. To examine the trade-off between sensitivity and specificity, we report the average ROC curves for classifiers in Table IV (see Fig. 5). Fig. 5 shows that the ROC curve for the XGB classifier trained using CCAFS integrated view, CCAFS_XGB, almost dominates all other ROC curves. In other words, for any choice of specificity between 0.95 and 0.4, CCAFS_XGB has a corresponding better sensitivity than the rest of the classifiers considered in this experiment.

**Figure 5.**
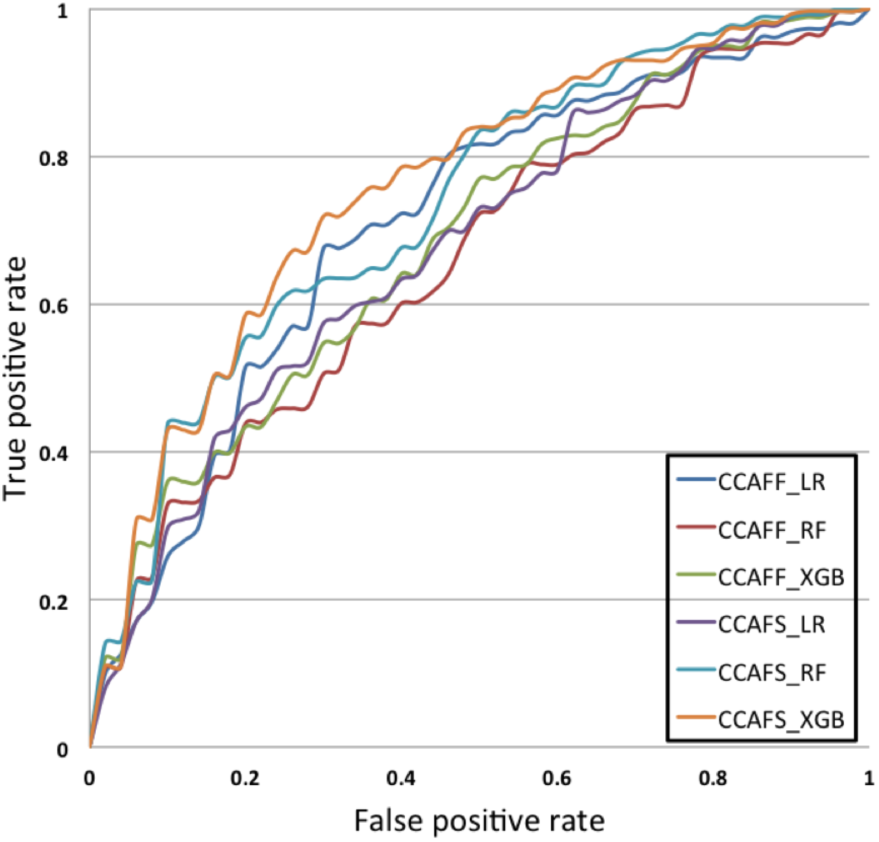
Average ROC curves for top performing CCAFF and CCAFS based models for predicting KIRC survival using three multi-omics views.

Next, we provide an analysis of the top multi-view features used by the best performing model with AUC equals 0.76.

### D. Analysis of Top Selected Multi-view Features

To get a robust estimate of feature importance, we evaluated the best model (XGB classifier using top 100 features selected using CCAFS with 50 CCA components) 100 times using different random partitioning of the data into 80% for training and 20% for test. In each run, the top 100 selected features were recorded and the feature importance scores of these features were incremented by 1. Table V summarizes the top 20 features selected from each view. We note that the average scores for top selected RNA-Seq, RPPA, and CNV are 0.45, 0.37, and 0.12, respectively. Interestingly, ranking the three views by their average feature importance scores is in agreement with ranking them by the highest AUC that could be reached using view-specific models (See Fig. 2).

**TABLE V.**
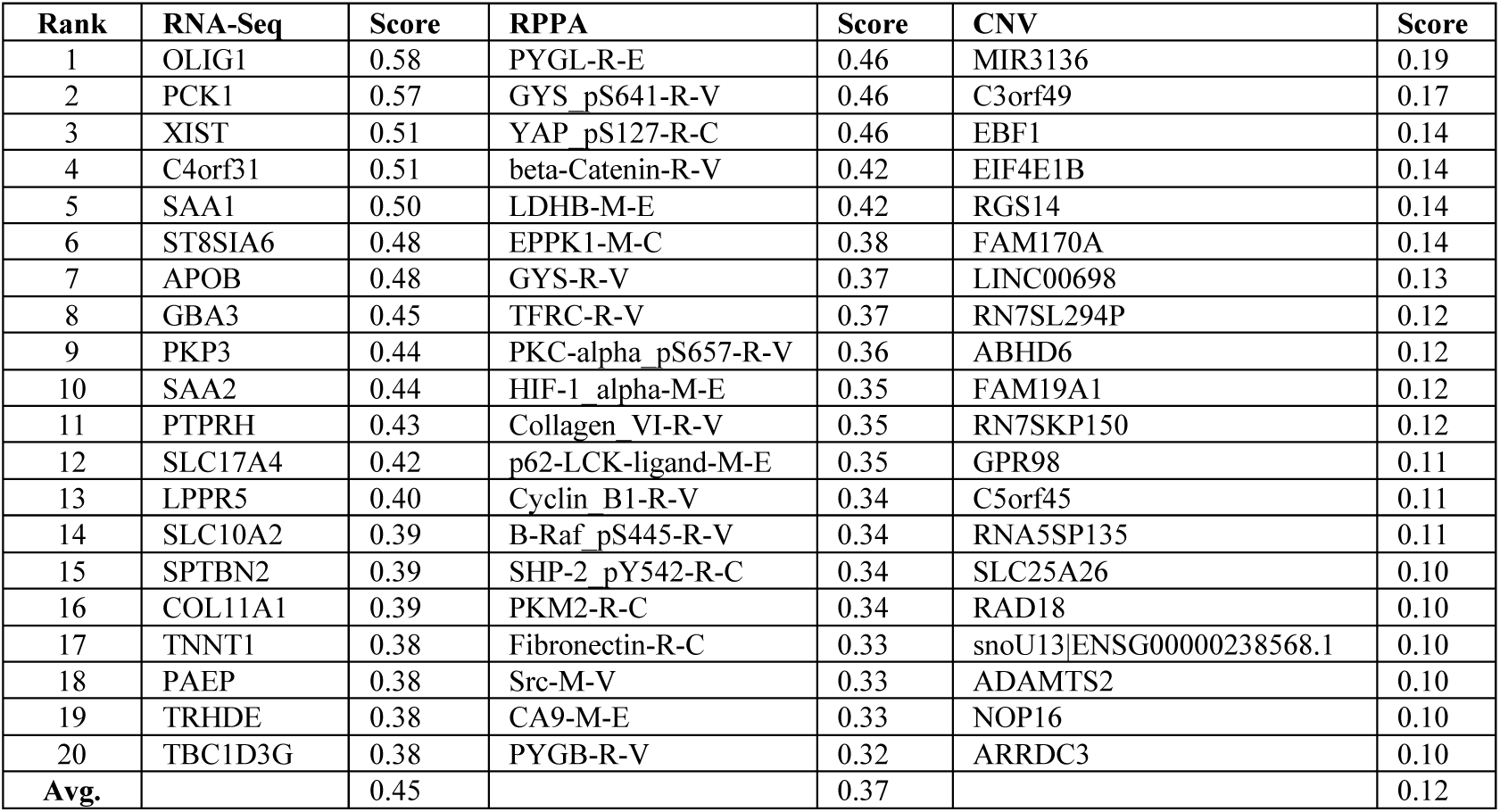
Top 20 features selected from each view and their importance scores. Reported scores are averaged over 100 runs of CCAFS method evaluated using 80% of the data for training and the remaining samples for test.

Similarly, we examined the best view-specific model (LR classifier using top 10 features selected using Two-Stage RF-based embedded filter). Table VI summarizes the top 10 selected RNA-Seq features. Surprisingly, only 5 features (highlighted in bold in Table VI) out of the 10 RNA-Seq features are shared with the top 20 RNA-Seq features selected using CCAFS method. This observation suggests that selecting features jointly from the three views is more likely to yield a set of features that are different from those selected independently from each single view.

**TABLE VI.** Top 10 RNA-Seq features selected using 100 runs of Two-Stage RF-based embedded filter. Bold text highlights features overlapping with those selected using CCAFS method.

| Rank | RNA-Seq | Score |
| --- | --- | --- |
| 1 | CHST9 | 0.99 |
| 2 | **XIST** | 0.80 |
| 3 | **OLIG1** | 0.62 |
| 4 | **LPPR5** | 0.61 |
| 5 | **C4orf31** | 0.59 |
| 6 | PRAP1 | 0.47 |
| 7 | **PCK1** | 0.39 |
| 8 | AKR1B10 | 0.39 |
| 9 | DNAJB13 | 0.38 |
| 10 | HABP2 | 0.38 |

Lastly, we applied gene-set enrichment analysis to identify overrepresented gene ontology (GO) terms in the two sets of RNA-Seq genes in Tables V and VI. Specifically, we used the gene-batch tool in GOEAST (Gene Ontology Enrichment Analysis Software Toolkit) [35] to identify significantly overrepresented Biological Processes GO terms for the two sets of RNA-Seq genes. We also used the Multi-GOEAST tool to compare the results of the enrichment analysis of these two sets. We noted that the set of 20 RNA-Seq genes identified by our multi-view feature selection method was enriched with only 9 GO terms while the smaller set of 10 RNA-Seq genes determined using single-view feature selection method was enriched with 20 GO terms. Four statistically significant (*p*-value less than 0.0003) overrepresented GO terms related to carbohydrate biosynthetic process were in common between the two sets. The remaining GO terms enriched in the set of 10 genes belong to carbohydrate metabolic process, small molecule metabolic process, and single-organism biosynthetic process. Interestingly, two statistically significant overrepresented GO terms (acute inflammatory response, p-value = 1.15e-5) and (acute phase response, p-value = 1.45e-6) were found to be enriched only in the set of the 20 genes. An acute inflammatory response is known to be beneficial in response and tissue damage and could have a role in tumor suppression [36].

## IV. Conclusion

Integrative analysis of heterogeneous high-dimensional multi-omics data, such as somatic mutation, copy number alteration (CNA), DNA methylation, RNA-Seq gene expression, and protein expression, is a major challenge in translational bioinformatics. Multi-view feature selection is a promising approach for integrating multi-omics data in a manner that takes into consideration interactions within and/or between different omics data sources. In this work, we presented a novel multi-view feature selection algorithm based on the well-known canonical correlation analysis (CCA) statistical method. We demonstrated the effectiveness of our proposed method in developing *integrated models* for the challenging task of predicting kidney renal clear cell carcinoma (KIRC) survival.

A major limitation of CCA-based feature fusion/selection methods described in this study is that feature fusion/selection was performed using *unsupervised* CCA. We conjecture that incorporating the labels of training data could lead to learning a better common subspace. Our ongoing work aims at incorporating *supervised* variants of CCA method (e.g., [37, 38]) into our framework. Our future work will examine the utility of kernalized CCA methods for capturing nonlinear relationships among the views.

## Acknowledgment

Research supported in part by the Center for Big Data Analytics and Discovery Informatics at the Pennsylvania State University and by the National Center for Advancing Translational Sciences, National Institutes of Health, through Grant UL1 TR000127 and TR002014. The content is solely the responsibility of the authors and does not necessarily represent the official views of the NIH.

